# “Ghost ponds” – How to resurrect in-filled farmland ponds to assist aquatic biodiversity conservation in agricultural landscapes

**DOI:** 10.1101/831859

**Authors:** Emily Alderton, Carl D. Sayer, Jan C. Axmacher, Ian R. Patmore, Helene Burningham, Phill L. Brown, Geoff Nobes

## Abstract

Growing recognition of the importance of ponds for landscape-scale biodiversity has led to considerable interest in their conservation, focusing on new pond creation, or existing pond restoration. However, there is a third approach; the re-excavation of ‘ghost ponds’ – former ponds deliberately filled-in due to agricultural intensification. Previous work has shown ghost ponds to retain viable sediment propagules of many aquatic plants for over a century, allowing for the rapid re-colonisation of resurrected pond sites. Here we detail the practicalities of the ghost pond resurrection approach, describing how to locate, identify, and excavate ghost ponds in agricultural land. We also report on colonisation by aquatic macrophytes and water beetles (Coleoptera) for three ghost pond resurrections in Norfolk, eastern England and make comparisons with neighbouring extant ponds restored to open-canopy conditions via major scrub and sediment removal at the same time. Ecologically important macrophyte taxa, including charophyte and *Potamogeton* species, successfully established in the ghost ponds and within one year they supported a comparable species diversity to the adjacent restored ponds. Our findings show that, where appropriate to land management goals, ghost pond resurrection could be a very valuable conservation approach within farmed landscapes.

## INTRODUCTION

Farmland ponds afford crucial oases within intensively managed agricultural landscapes, contributing to habitat heterogeneity (Benton *et al.* 2003), and to aquatic and terrestrial biodiversity (Williams *et al.* 2003, Davies *et al.* 2016). Despite their ecological importance, over 50% of farmland ponds in the UK have been lost during the last century, with many filled in due to agricultural land consolidation (Hassall 2014). Similar patterns of pond loss have occurred across Europe, with severe negative consequences for aquatic biodiversity (Zacharias & Zamparas 2010, Curado *et al.* 2011).

Awareness of the importance of ponds for aquatic biodiversity conservation has grown in recent years (Oertli *et al.* 2018). Currently, new pond creation (Biggs *et al.* 2005, Williams *et al.* 2010) and the restoration and subsequent management of highly terrestrialised extant ponds via scrub and sediment removal (Sayer *et al.* 2013) are major pond conservation approaches utilised in the UK. Both approaches aim to create mosaics of ponds at different successional stages, and both have been shown to successfully increase landscape-scale aquatic biodiversity (Thiere *et al.* 2009, Sayer *et al.* 2012). Here we present a new approach in pond conservation; the re-excavation of ‘ghost ponds’ - ponds formerly filled in, especially during the 1930s-1970s, as part of a more general process of agricultural intensification and land consolidation.

Our recent greenhouse and mesocosm-based experiments have shown at least 9-10 aquatic macrophyte species to germinate from seeds and oospores, following 50-150 years of enforced dormancy in the sediments of ghost ponds (Alderton *et al.* 2017). The rapid (< 6 months) re-colonisation of resurrected ghost ponds by a diverse aquatic vegetation similarly suggests a strong seed-bank influence. Ghost ponds therefore represent abundant, hitherto little exploited time capsules of aquatic plant diversity in agricultural landscapes affording opportunities for enhancing landscape-scale aquatic biodiversity and connectivity.

In this paper we provide details on how to locate, identify and excavate ghost ponds, illustrated with examples of three ghost pond resurrections undertaken in the villages of Westfield (Pond GP_150_), Guestwick (Pond GP_50_) and Stody (Pond GP_45_) in Norfolk, eastern England. In addition, we highlight the short-term (1 year) response of macrophytes and water beetles (Coleoptera) to these ghost pond resurrections and compare the results to three neighbouring (within 250 m) extant ponds (ponds WERE, GURE, STRE paired with GP_150_, GP_50_ GP_45_ respectively) restored top open-canopy conditions via major scrub and sediment removal at the same time. Norfolk is a dry (average 600-700 mm rainfall per annum), predominantly arable agricultural region, which has experienced pond losses of around 28% since the early 1950s (30,088 ponds mapped on 1953-1957 OS maps, compared to 21,697 ponds on the 2014 OS map - Alderton 2017), with the majority of remaining ponds being heavily terrestrialised, rendering them of relatively low conservation value (Sayer *et al.* 2012). While these pond losses are low compared to other UK regions (Wood *et al.* 2003, Hassall 2014), they nevertheless represents a considerable reduction in pond numbers, and hence ghost pond resurrection aimed at increasing the number of open-canopy ponds in the landscape could make a substantial contribution to farmland biodiversity conservation.

## METHODS

### Locating ghost ponds

The first step in ghost pond resurrection is to locate lost, in-filled ponds on historic maps. Ghost ponds can usually be easily located by overlaying recent maps with different editions of historic maps, satellite imagery and aerial photography (**Figure 1**). Further, there is also much to be gained by talking to local people, especially the farm workers and land managers who were sometimes actually involved with ponds being in-filling. In the case study presented here, we obtained recent and historic (19^th^ and 20^th^ Century) Ordinance Survey (OS) maps (EDINA Digimap 2013), 18^th^ and 19^th^ century tithe maps (Norfolk County Council 2012), and satellite imagery from Google Earth for the pond areas. Maps and imagery were georeferenced in QGIS (QGIS Development Team 2009) to British National Grid for geospatial comparison. Using a combination of historic maps from different time intervals allowed for estimates of ‘time since burial’ for each ghost pond, based on the most recent map demarcation of a pond. Overlaying historic maps with modern satellite imagery is particularly helpful for locating ghost ponds in the field. In the case of the three ghost ponds studied here, GP_150_ was filled-in during the mid-1800s (pond marked on 1839 tithe map, but not on the 1883 OS map), GP_50_ was filled-in during the late 1960s and GP_45_ was filled-in during the early 1970s.

**Figure 1.**
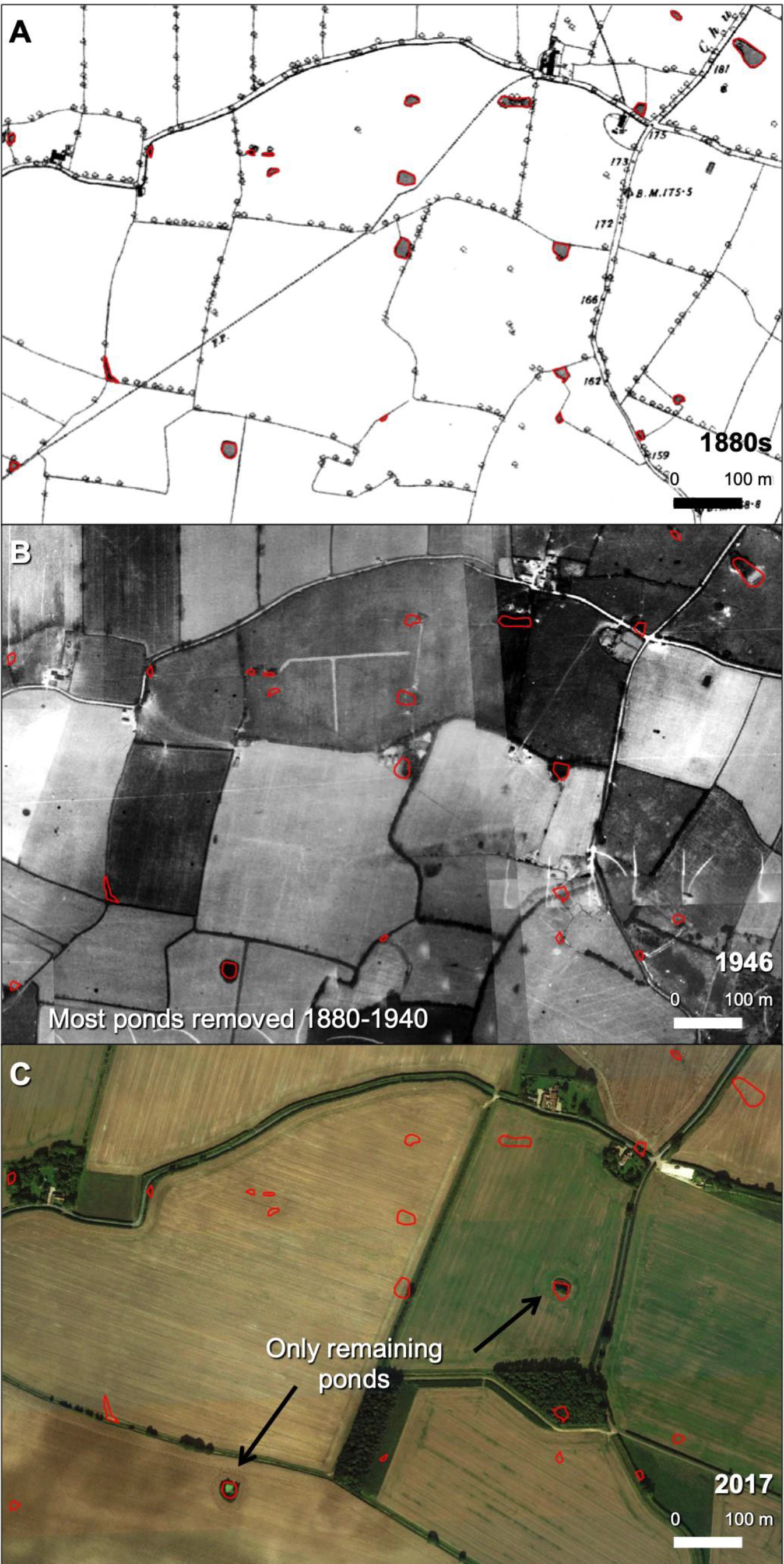
Dramatic pond loss illustrated for an area of mid-Norfolk by comparing ponds present in the 1880s (a), in 1946 (b) and in 2017 (c). Ghost Ponds present in 1946 and 2017 are marked by red empty polygons.

After locating ghost ponds on the map, a field visit should be conducted to confirm their exact location on the ground. Ghost ponds commonly remain visible as damp depressions in fields (particularly during winter when a low crop cover coincides with high water tables – **Figure 2a,b**), or as crop marks due to differing soil moisture content and quality and hence differing rates of crop maturation (**Figure 2c**). Some ghost ponds, however, may only be indicated by slight differences in soil colour, or by a barely evident topographic low (**Figure 2d**). In cases where there is little or no visible evidence for a ghost pond on the ground, but a pond is clearly marked on historic maps, topographical data available from Lidar surveys or acquired locally using GPS or unmanned aerial systems (UAS) can be useful in detecting subtle changes in elevation (**Figure 3**).

**Figure 2.**
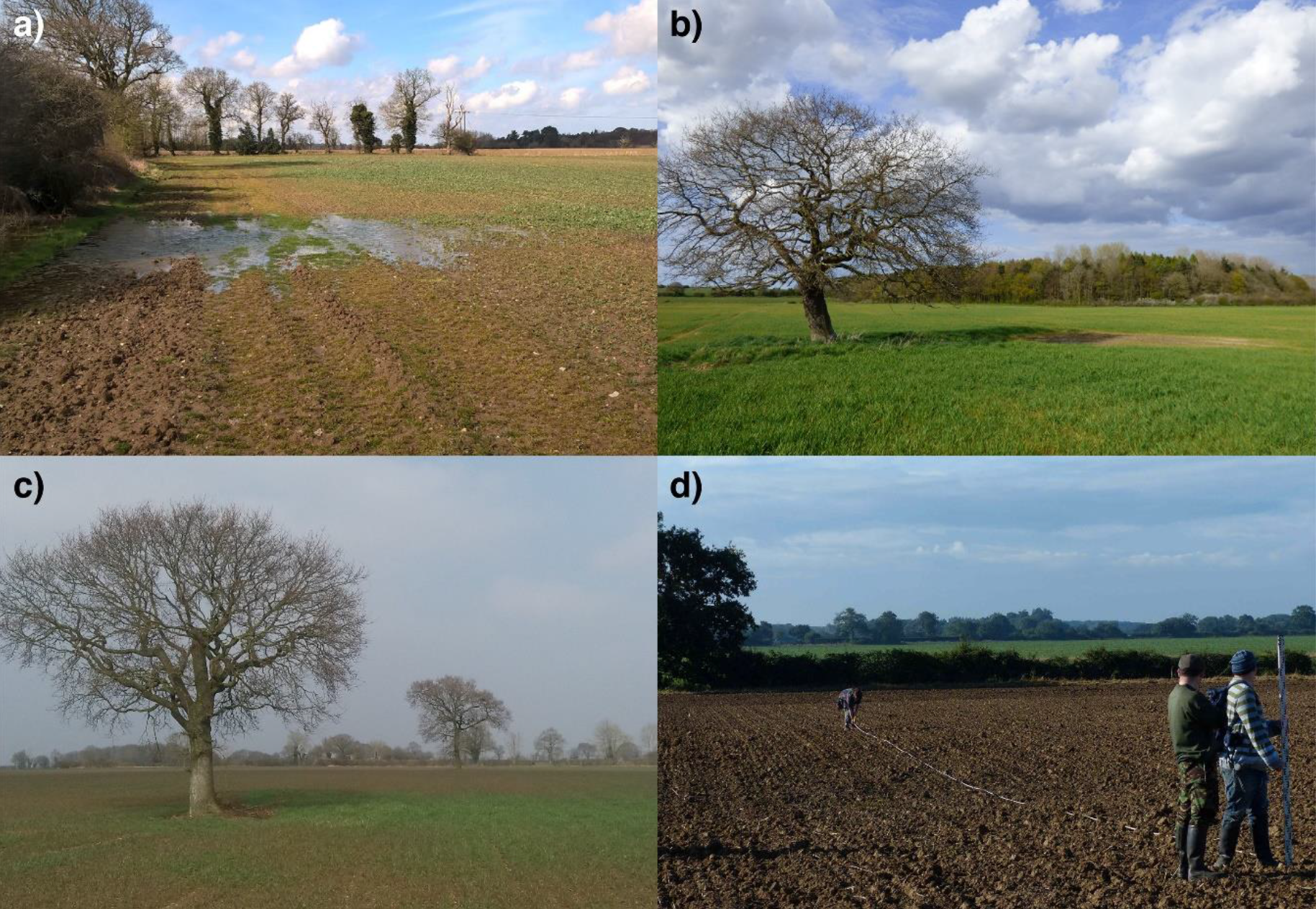
Ghost ponds are often visible in the fields as damp depressions which often flood after heavy rain (a), as patches of poor crop cover (b) as crop marks where the crop matures at a different rate compared to the surrounding area (c) or as subtle topographic lows in a field (d).

**Figure 3.**
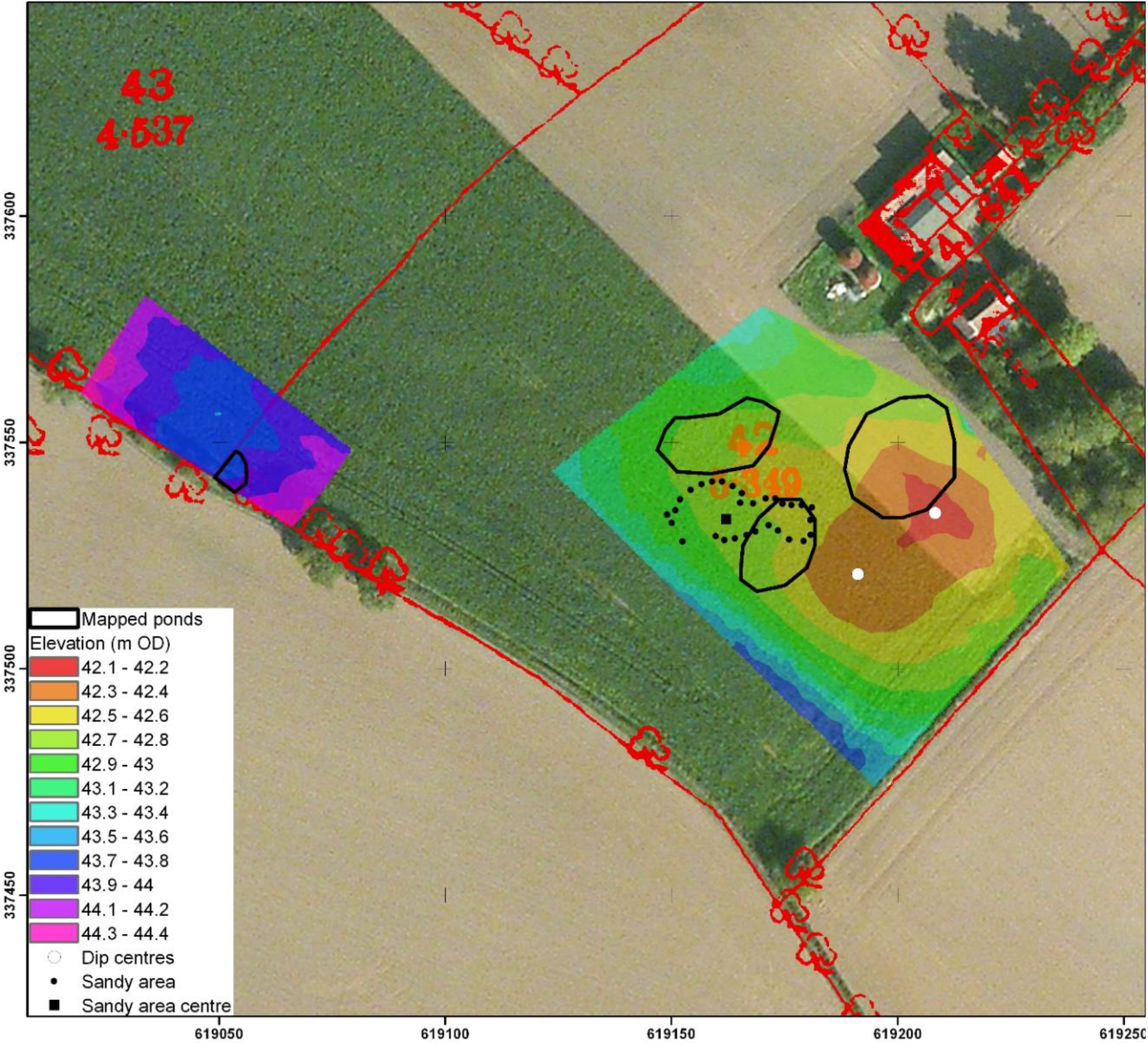
Digital Elevation Model (DEM) of four ghost ponds in North Norfolk, eastern England, marked on the 1^st^ Edition OS map (1886), but absent from later editions. DEM overlays satellite imagery and 1886 OS map outlines.

In all cases, it is important to recognize that ‘pond-indicating features’ may not exactly match the location of a ghost pond as defined on the historic maps. For example, **Figure 3** shows a survey of four ghost ponds in an arable field where ponds evident on the 1885 OS map are compared with a Digital Elevation Model (DEM) generated from a high-resolution dGPS survey. A clear mismatch between surface topography and mapped pond location is evident which could be explained by inaccuracies in historic mapping or changes in surface topography since pond burial. If the ponds were filled by scraping top-soil from the immediate surroundings into the pond, this could alter the size and shape of the depression around the original pond basin. Activities such as ploughing could further change the shape of the depression, as the wet pond sediment and overlying soil compact over time. It is also worth noting that not all depressions or hollows appearing in agricultural fields are ghost ponds; dry pits, collapsed field drains, or natural changes in elevation also occur. Hence, a multi-angled approach considering different editions of historic maps alongside other geospatial data such as surface topography is often necessary to the search for a ghost pond.

Most ghost ponds can be accurately located on the ground using the combination of recent and historic maps, combined with field measurements (distance from hedgerows or other mapped features), and visual inspection of vegetation, soil colour and topographical changes. In cases where the location of a ghost pond remains unclear, determining soil stratigraphic profiles at intervals across the region of interest can sometimes be useful to ascertain the location and extent of the ghost pond. Specifically, obtaining stratigraphic evidence using a soil auger or by excavating a small, vertical-walled pit can be crucial in understanding the nature of infilling and location of the pond relative to the agricultural surface. The pond infill visually comprises a mix of local soil (backfill) and agricultural waste providing a clear boundary with the original pond bed at depth.

### Excavating ghost ponds

Once ghost ponds have been located (**Figure 4a**), excavation can begin. Ghost ponds should ideally be resurrected post-harvest or during fallow field years, during the autumn to early winter months. At this time of year the water table is low, affording suitable conditions for working with heavy machinery and for inspecting sediment stratigraphy during the early stages of excavation.

**Figure 4.**
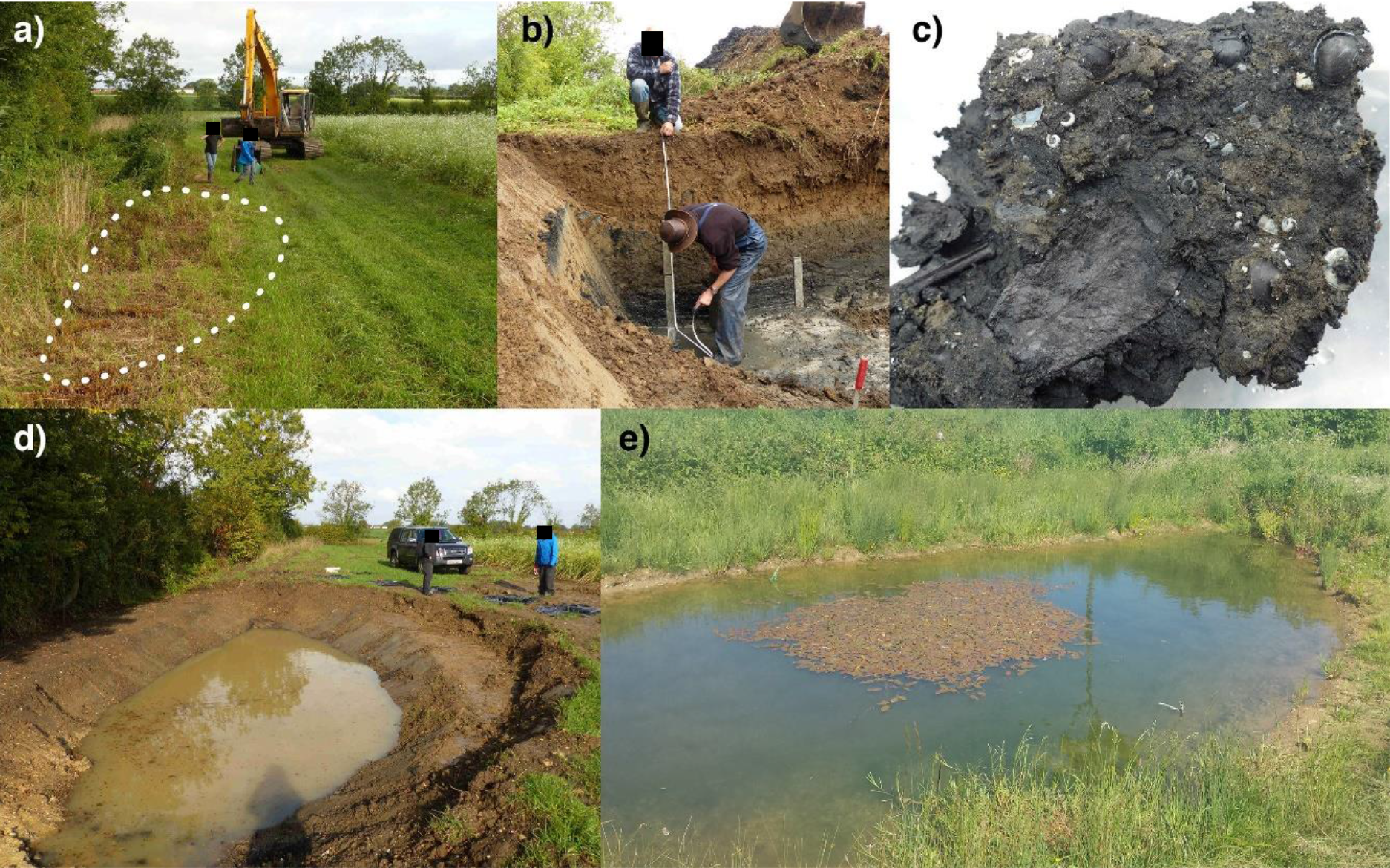
Excavation and monitoring of pond GP_150_. a) GP_150_ prior to excavation with ghost pond outline highlighted by white dots; b) Sampling the historic sediment layer in the test trench; c) Historic pond sediment containing sub-fossil remains of Mollusca; d) GP_150_ in October 2013, three weeks after excavation; e) GP_150_ in May 2015 twenty months after excavation.

In excavating a ghost pond, it is important to stay faithful to its history and hence, as far as possible, excavate the pond to its original dimensions. Hence, the first stage in ghost pond excavation is to establish the depth and extent of the historic pond sediment layer, which can be used to approximate pond shape and size. This can be achieved by digging a test trench through the suspected centre of the ghost pond (based on map demarcation and field measurements), until the historic pond sediment layer is reached (**Figure 4b**). Subsequently, excavation of a second trench located at right angles to the first (forming a cross), is also useful to help gain knowledge of the ponds former dimensions. A 14-tonne 360 degree digger (with an experienced driver), is highly suitable for excavating both the test trench and the ghost pond itself. Pond sediment typically consist of fine, black, silt, easily distinguished from the coarser, lighter top soil used to fill-in ghost ponds (**Figure 4b**). This sediment may also contain easily visible preserved remains of aquatic plants (especially whole leaves and seeds) and animals, such as Mollusca (**Figure 4c**). The historic sediment layer is an important source of aquatic propagules (Alderton *et al.* 2017), and it is important that at least part of this sediment remains after excavation. Sediment samples for propagule and palaeoecological analysis can be collected using a box core from exposed pond sections (**Figure 4b**).

Working outwards from the test trench(es) with the digger, the pond should be excavated following the contours of the historic sediment profile, so that the resurrected pond has a bathymetry that matches, as far as possible, the former bathymetry of the ghost pond. Care should be taken not to remove too much of the past pond sediment by excavating a shallow layer of top soil at a time. Careful excavation also allows for the retrieval of potentially informative discarded objects often thrown into ponds before and during burial. Old glass bottles, clay pipes, and even plastic food packaging (as was the case for pond GP_50_ where bags for frozen peas were found), can provide dateable evidence for the approximate time of burial of a pond. Thus it is important to record any significant buried finds during the excavation process. In addition, given that in-filled ponds often resulted in wet patches in fields, many farmers subsequently installed field drains at these locations and it is important to suitably remove or block these drains during excavation such that water freely enters the pond and importantly does not quickly drain out of it.

Once excavation is complete, a margin of at least 7-10 m width should be marked out around the pond as a buffer from farmland activities and agro-chemical applications. In this respect it is important to have an agreement in place with the farmer before any works take place. Newly excavated ghost ponds can be left to fill naturally with rainwater through the winter, and importantly, both the pond and its margin should be left to colonise naturally. For our three study sites, excavation took less than one day per site, with a professional digger driver working alongside 1-2 people on the ground who guided the driver by helping to identify when the digger bucket had reached the historic pond sediments. All study sites were excavated in September-October 2013.

### Pond restoration monitoring

Our three ghost ponds and the three paired restored ponds were closely monitored throughout the first year of colonisation (Year 1: September 2013-September 2014). During year one, aquatic macrophyte surveys were conducted in October-November 2013 (week 5), January 2014 (Week 12), February 2014 (week 16), May 2014 (week 28), June 2014 (week 34), July 2014 (week 40), August 2014 (week 44), and September 2014 (week 50). All wetland-associated plants were recorded visually with the assistance of a double-headed rake. Water beetles (Coleoptera) were surveyed in May 2014 and September 2014 as a proxy for broader invertebrate diversity (Briers & Biggs 2003), and because they are regarded as some of the earliest colonists of newly excavated ponds (Coccia *et al.* 2016). Each pond was sampled exhaustively for water beetles using a 1 mm mesh pond net swept through the different available meso-habitats until no new species were encountered. If possible, all adult water beetles were identified to species-level in the field and released back into the pond. Where individuals could not be safely identified at the pond they were preserved in 70% industrial methylated spirits (IMS) and identified at a later stage with the assistance of a dissecting microscope.

## RESULTS

### Wetland macrophytes

Prior to excavation, the three ghost pond sites provided little ecologically valuable habitat: the agricultural field containing GP_45_ had not been cultivated in 2013 due to heavily water-logged soil, and supported a low diversity of common agricultural weeds (mostly stinging nettle *Urtica dioica*). The site of GP_50_ had been cultivated, harvested and ploughed, and contained no vegetation. Due to its location within a topographic low at the field margin pond GP_150_ contained some common wetland species including soft rush *Juncus effusus*, hard rush *Juncus inflexus*, hemp-agrimony *Eupatorium cannabinum*), and a mix of common agricultural grasses in dryer areas. All these species rapidly returned to the site after pond excavation.

Macrophyte colonisation of the ghost ponds is illustrated in **Figure 5**. The ghosts filled with water over winter and the first submerged aquatic species to colonise the newly excavated ghost ponds were common water-crowfoot *Ranunculus aquatilis* (GP_150_) and hairlike pondweed *Potamogeton trichoides* (GP_45_), both appearing by January 2014 (week 12). Dense beds of common stonewort *Chara vulgaris* (later joined by opposite stonewort *Chara contraria* and fragile stonewort *Chara globularis*) began to develop in GP_50_ and GP_150_ by May 2014 (week 28). In GP_45_, charophyte beds developed later (by July 2014, week 40), with a mixed initial assemblage of *C. vulgaris, C. globularis*, and bristly stonewort *Chara hispida*. Broad-leaved pondweed *Potamogeton natans* began to appear between weeks 34 (GP_150_) and 40 (GP_45_ and GP_50_), followed later in the year by curled pondweed *Potamogeton crispus* and horned pondweed *Zannichellia palustris* (weeks 40 and 44 respectively).

**Figure 5.**
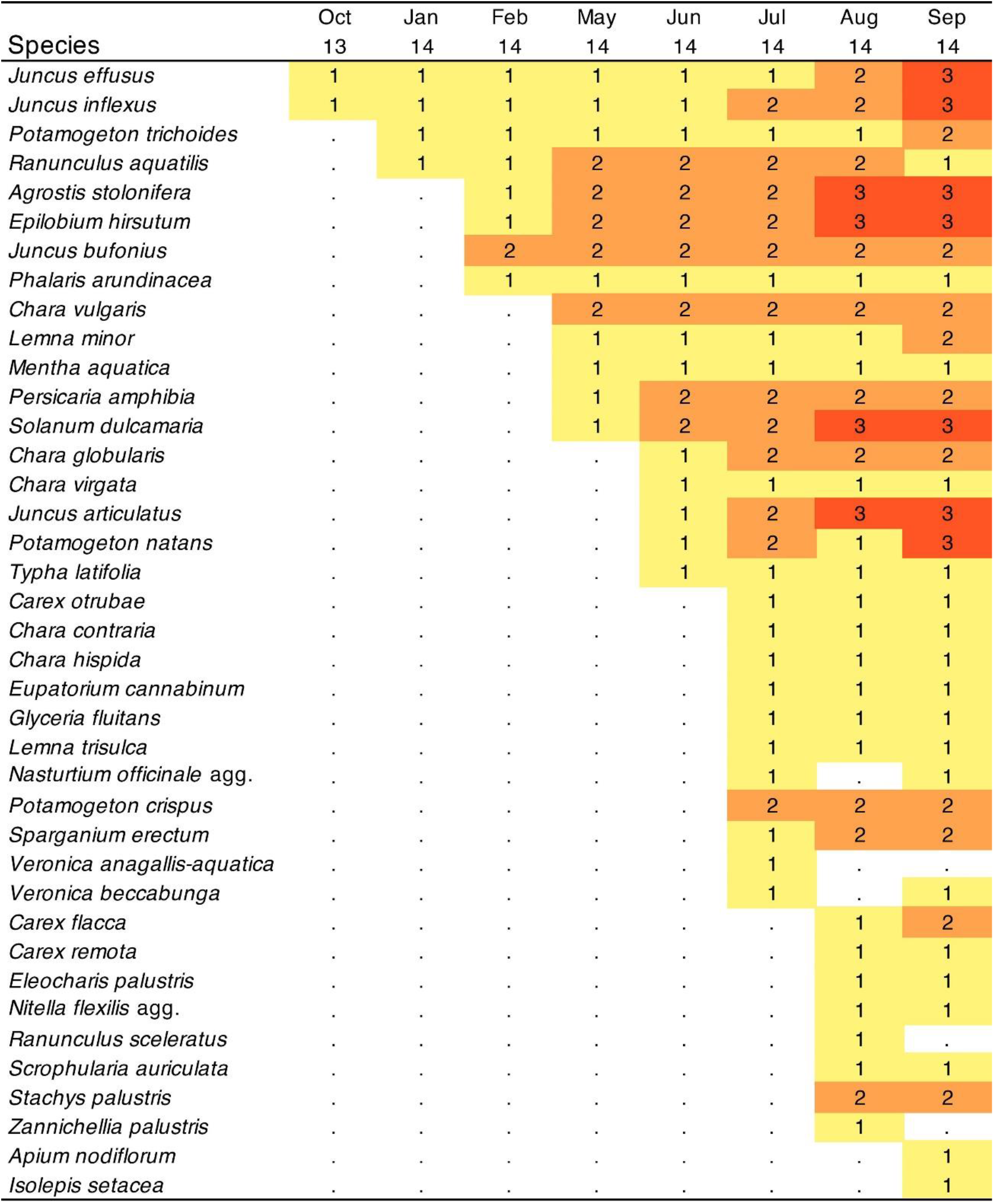
Accumulation of wetland macrophyte species in study ponds GP_45_, GP_150_, GP_50_ and Coloured highlight indicates whether plants occurred in 1 (yellow), 2 (orange) or all 3 (red) of the study ponds. Species which also germinated in controlled experimental conditions (see Alderton *et al.* 2017) from at least one of the three study ponds were: *Nitella flexilis* agg. *Potamogeton natans*, *Chara contraria*, *Chara hispida, Chara globularis*, *Chara vulgaris*, *Chara virgata*, *Ranunculus aquatilis*, *Juncus* sp.

Despite the rapid colonisation of aquatic macrophytes, all three ghost ponds remained in an algal-dominated state for the first 8-9 months, with mostly bare or crop-dominated pond margins generated by the persisting crop seedbanks. However, the initially poor appearance of the sites was no cause for concern; after one year, all three ghost ponds had clear waters and supported a high abundance and diversity of wetland macrophytes, while the surrounding margins supported a range of native wildflower species including primrose *Primula vulgaris*, bee orchid *Ophrys apifera*, field forget-me-not *Myosotis arvensis*, purple loosetrife *Lythrum salicaria*, gypsywort *Lycopus europaeus*, and meadow buttercup *Ranunculus acris*.

The importance of the historic propagule bank for ghost pond colonisation was evident, with at least eight macrophyte species recorded in the study ponds also found during viability tests of the historic propagule bank (Alderton *et al.* 2017). Other species may also have originated from historic pond sediments, but could have been missed in the viability tests due to the small sample sizes used in the tests or to sub-optimal germination conditions for some species in the experimental settings.

Overall, the resurrected ponds provided habitats of moderate to high wetland plant assemblage conservation value (Biggs *et al.* 2005), with GP_45_, GP_50_ and GP_150_ supporting 25, 18 and 27 species respectively, at the end of their first year (**Figure 5**). Compared to typical UK lowland ponds (0-27 species, average 12 species; Williams *et al.* 1998), the number of recorded plants in the ghost ponds was also relatively high, whilst the rate of species accumulation and species number after one year of colonisation was similar to that observed at the neighbouring restored ponds (**Figure 6a**).

**Figure 6.**
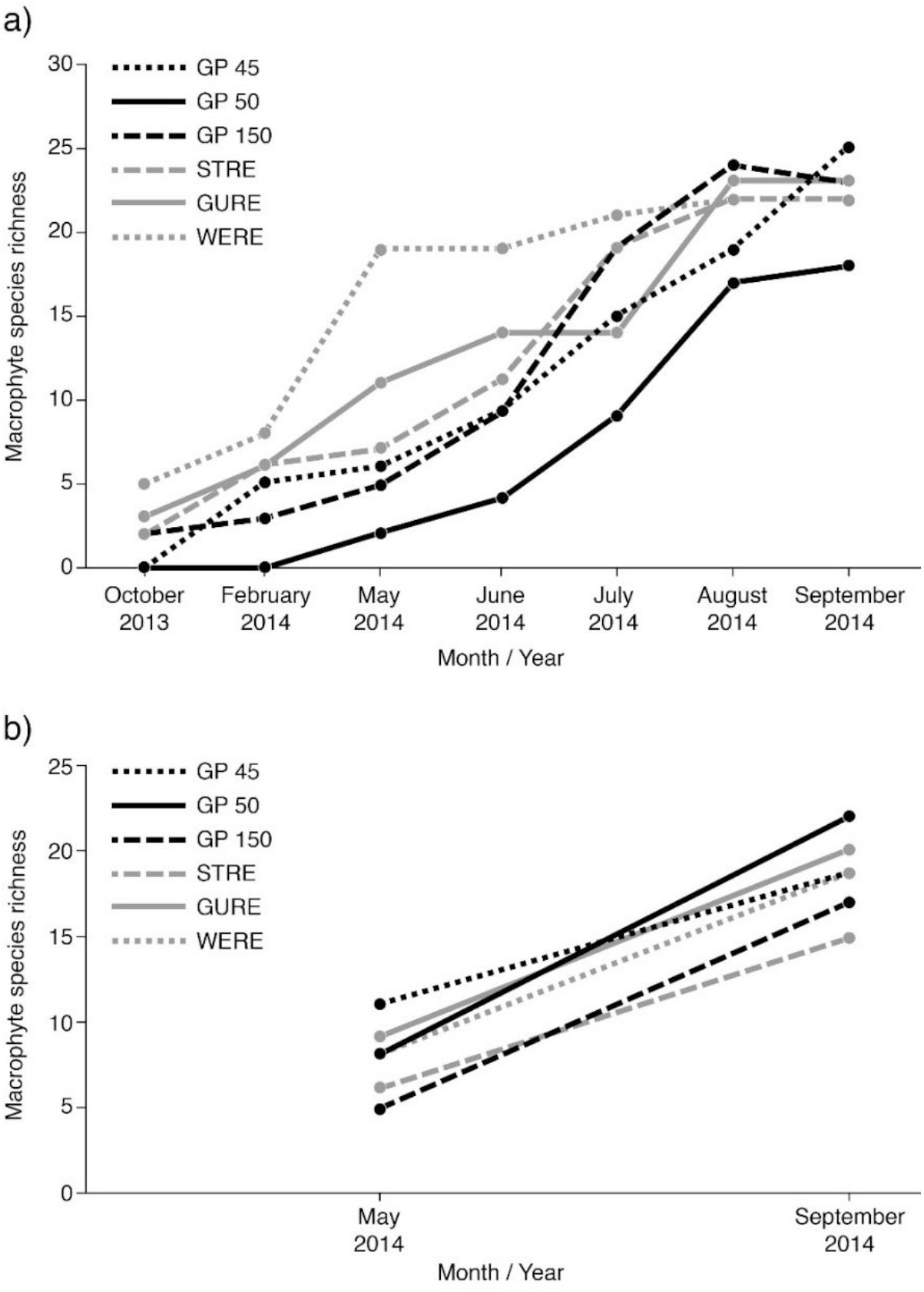
Accumulation of wetland plants (a) and water beetles (b) in the resurrected ghost ponds (GP 45, GP 50, GP150) and adjacent ponds restored by major scrub and sediment removal (STRE, GURE, WERE).

### Water beetles

The three ghost ponds were rapidly colonised by water beetles, with a total of 35 species (representing six families) recorded in year one (**Table 1**). Species arrivals strongly reflected known seasonal flying times (Miguélez & Valladares 2008, Boda & Csabai 2013) and it was notable that, in all ponds, species richness increased between the surveys undertaken eight and twelve months post-resurrection. Individual study ponds contained between 20 (GP_50_ & GP_150_) and 24 (GP_45_) water beetle species, with the shallow and sandy GP_45_ supporting the highest number of unique species (n = 8). Average species richness in the ghost ponds was comparable to that observed for other open-canopy restored farmland ponds in Norfolk sampled in a comparable manner (16-19 species across 4 ponds; Law *et al.* accepted). Further, as for wetland plants, the accumulation and final number of water beetles in the ghost ponds was similar to that observed for the nearby extant ponds restored by scrub and sediment removal (**Figure 6b**).

**Table 1.**
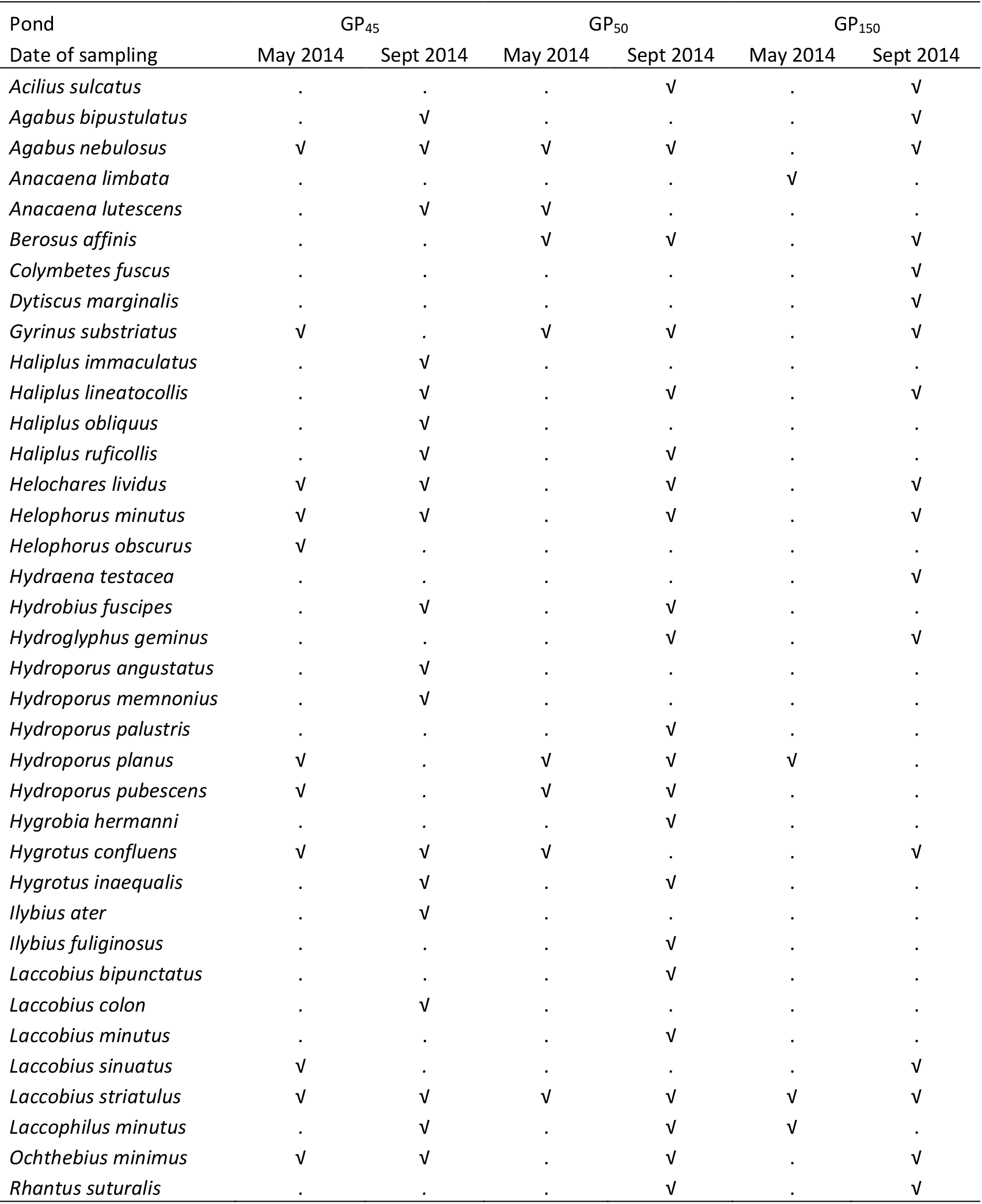
Water beetles colonising the three studied ghost ponds during Year 1. The May and September samples were collected eight (May 2014) and twelve (September 2014) months following ghost pond resurrection respectively.

## DISCUSSION

The resurrection of ghost ponds clearly represents an effective means of restoring high-quality pond habitats in agricultural landscapes. Ghost ponds are easily identified on maps and are often relatively straightforward to locate in the field, and can be excavated cheaply and quickly within just one day. Since ghost ponds generally represent sub-optimal patches of farmland due to high water-table positions and poor drainage within their footprint, excavation will have negligible impacts on crop yields. This is especially true of grass headlands at field edges (as for GP_150_ – **Figure 4a**) and permanent grassland or woodland, where the resurrection of a pond interferes minimally with agricultural activities.

Even after burial for over 150 years, the historic propagule bank provides a source of native macrophyte species, which rapidly re-colonise and fill ghost pond basins (**Figure 4e**). We found that, within just one year, agricultural ghost ponds reached comparable levels of macrophyte and aquatic beetle diversity to local restored ponds and to existing high quality lowland ponds both in the study region. We suggest that rapid colonisation from the historic propagule bank made an important contribution to early macrophyte diversity in the ghost ponds, although more work, particularly direct comparisons between colonisation processes in ghost ponds and new ponds, is needed to prove this hypothesis. While propagule banks are known to be important for species ‘dispersal through time’ in temporary wetlands (Britton & Brock 1994), importantly, our findings suggest the same principles apply to buried farmland ghost ponds.

This study clearly detected a number of common species in the historic propagule bank that represented a range of genera and seed types, suggesting that many more species might be able to survive prolonged burial underneath agricultural fields. This opens up the possibility that ghost ponds could act as ‘time-capsules’ for locally scarce or even locally extinct macrophyte species or of otherwise extinct local ecotypes, thus contributing to aquatic biodiversity conservation at both species and genetic levels. Indeed, a number of studies have documented the re-appearance of rare or locally extinct species from historic propagule banks (Weyembergh *et al.* 2004, Scott *et al.* 2012, Kaplan *et al.* 2014), but it has been generally assumed that this would not be possible in arable agricultural settings. Our study provides evidence to the contrary and also reveal propagule longevity to be on much longer (centennial) timescales than previously thought. Clearly more research is required on propagule longevity for a larger sample of farmland ghost ponds and aquatic plant species. In addition, more work is now needed to test the potential and success of ghost pond revival covering a broader range of pond types (e.g. dew ponds, natural pingo ponds) as well as in different biogeographical, geological and hydrological settings.

Given the high abundance of ghost ponds across many of the UK’s and indeed European farmland landscapes, combined with the speed, cheapness and relative simplicity, with which these ponds can be resurrected, we urge conservationists to incorporate ghost pond resurrection into landscape-scale conservation planning and strategies. Ghost pond resurrection may be an extremely important tool for enhancing aquatic connectivity in fragmented aquatic landscapes and should be considered on an equal footing to pond creation and restoration. Equally, we advocate that agri-environment schemes need to much more fully embrace the ghost pond resurrection approach to maximise the funded pond conservation options available to farmers. Not only is ghost pond resurrection of great importance to conservation, but it is inspiring for conservation practitioners and nature-loving farmers too, as it involves turning back the clock and thinking about and reconstructing farmland pond landscapes of old. When you see a mysterious “*will-o’-the-wisp*” in a field or a dark patch in the plough soil in the falling light of winter, realise that a pond ghost lies beneath, buried alive and ready to spring to life again, if such is wished for.

## ACKNOWLEDGEMENTS

This research was funded by a Natural Environment Research Council (NERC) PhD award to Emily Alderton (Ref: NE/K501037/1), with additional funding kindly provided by the Norfolk Biodiversity Partnership, ENSIS Ltd., and the UCL Graduate School. We thank some brilliant landowners who made the study possible: J.D & N.J Anema, M.B & R.A Jensen Farming, and the Stody Estate Ltd. (especially Ross Haddow). We also thank Dominic Arnold for expertly working the digger, Derek Sayer for much logistical support and Emily Smith, Phoebe Lewis, Ben Siggery, Arnie Warsop and Bernard Cooper for field assistance.

